# SepN is essential for assembly and gating of septal junctions in *Nostoc* sp. PCC 7120

**DOI:** 10.1101/2022.01.26.477872

**Authors:** Ann-Katrin Kieninger, Piotr Tokarz, Gregor L. Weiss, Martin Pilhofer, Iris Maldener

## Abstract

The multicellular life style of filamentous cyanobacteria like *Nostoc* sp. PCC7120 relies on a cell-cell communication system involving so called septal junctions. These are multiprotein complexes, which traverse the septal peptidoglycan through nanopores, connecting the neighboring cells and enabling molecule transfer along the filament. The intercellular communication is crucial when different cell types in the filament, vegetative cells and heterocysts, have to exchange metabolites and signaling molecules. Septal junctions of cyanobacteria can even control the molecule exchange by gating. The multiprotein complex consists of three modules: the septum spanning tube, the plug residing within the cytoplasmic membrane at both ends of the tube and a membrane associated cap module, covering the plug/tube modules on the cytoplasmic side of each neighboring cell. Until now, FraD was the only identified protein component of the septal junction protein complexes and localizes to the plug module. Here, we identified SepN as a new component via co-immunoprecipitation using FraD as bait and further demonstrated its essential role in septal junction assembly. Despite normal septal nanopore formation, a mutant in *sepN* exhibited a highly reduced rate of intercellular communication and was unable to gate the exchange of molecules. Cryo-electron tomography of cryo-focused ion beam thinned *sepN*-mutant filaments revealed septal junctions lacking the plug module and lateral cap openings. The combination of missing plug but present cap allowed to deduce the importance of the plug module in ensuring the correct cap architecture and, more importantly, in sealing the diffusion area in the closed septal junction state.

## Introduction

Multicellular organisms rely on cell-cell communication for proper growth and survival. Plants use plasmodesmata and metazoans gap junctions to exchange molecules intercellularly. A multicellular lifestyle in Bacteria evolved in filamentous cyanobacteria that perform cell differentiation under specific environmental conditions. Among these, *Nostoc* sp. PCC 7120 (also known as *Anabaena* sp. PCC 7120; hereafter *Nostoc*) is a model organism for cell-cell communication, a process that is enabled by septal junctions (SJs) (Kieninger & Maldener 2021).

Cyanobacteria are gram-negative bacteria, having an outer membrane, which in case of *Nostoc* and related species, encompasses the entire filament (Flores et al 2006) without entering the septum after cell division (Flores et al 2006). The peptidoglycan represents a contiguous giant macro molecule, which is not split inside the septum, (Hoiczyk & Baumeister 1995, Lehner et al 2013). Molecules determined to move between the cells of the filament have to pass the cytoplasmic membrane of each cell and the septal peptidoglycan.

Some years ago, the semi-regularly drilled perforations of 20 nm in diameter were discovered in purified septal PG discs from *Nostoc punctiforme* and *Nostoc* and named the nanopore array (Lehner et al 2013, Nürnberg et al 2015). Cell wall amidases are responsible for drilling these 50-150 nanopores per septal disk, which are essential for intercellular communication (Bornikoel et al 2017, Lehner et al 2011) (reviewed in: Kieninger et al 2019, Maldener & Forchhammer 2015). Molecular exchange in filamentous cyanobacteria is a simple diffusion event and can be traced using fluorescent dyes in fluorescence recovery after photobleaching (FRAP) experiments (Mullineaux et al 2008, Nieves-Morión et al 2017b). Previously described cell-cell connections (Giddings & Staehelin 1978, Lang & Fay 1971) were suggested to traverse these nanopores and directly link the cytoplasm of neighboring cells (Flores et al 2016, Mariscal 2014). These cell-cell joining structures most probably are of proteinaceous nature (Omairi-Nasser et al 2014, Wilk et al 2011) and were referred to as septal junctions (Flores et al 2016, Mariscal 2014). Recently, we revealed the *in situ* architecture of septal junctions in *Nostoc* by imaging focused ion beam (FIB)-thinned filaments with cryo-electron tomography (Weiss et al 2019). The septal junctions exhibited a five-fold symmetric cytoplasmic cap module, a cytoplasmic membrane-embedded plug domain and a tube flexible in length that traverses the septal PG connecting to the cap/plug module of the opposite cell (Weiss et al 2019). The septal localized protein FraD (Merino-Puerto et al. 2010) was shown to be a structural SJ component, which localizes to the plug domain (Weiss et al 2019). Mutants in other septal proteins, AmiC1, FraC, SepJ, SjcF1, SepI and GlsC have a similar phenotype of impaired nanopore array, reduced rate of molecular exchange, filament fragmentation (except for AmiC1 and SjcF1) and inability of diazotrophic growth (Berendt et al 2012, Bornikoel et al 2017, Flores et al 2007, Merino-Puerto et al 2010, Merino-Puerto et al 2011, Nieves-Morión et al 2017a, Rudolf et al 2015, Springstein et al 2020). However, mutants lacking SepJ, SjcF1, AmiC1 or FraC exhibited no altered SJ structure, and therefore are most likely no structural SJ components (Weiss et al 2019).

Interconnection of all cells within a filament also constitutes a risk for the organism when cells are attacked by predators (Bauer & Forchhammer 2021), burst due to shear forces (Nicolaisen et al 2009) or die of senescence. In such cases isolation of the injured cell by abolishment of molecular exchange seems to be essential for ensuring the survival of the remaining filament. Indeed, interrupted communication between senescent, terminally differentiated heterocysts (cells specialized for nitrogen fixation) and vegetative cells was observed (Nürnberg et al 2015). Furthermore, the cap structure undergoes a conformational change, which could lead to a closed state accompanied by abolished intercellular exchange, when facing stress conditions like disruption of the proton motive force, oxidative stress or prolonged darkness (Weiss et al 2019). The impact of the cap and plug module on SJ closure was further demonstrated by a *fraD* mutant, which lacked the cap and plug modules and was unable to abolish communication when exposed to stress conditions (Weiss et al 2019). Interestingly, the cap can switch back to the open, communication-allowing state when conditions are more favorable, which renders SJs gated cell-cell connections functionally analogous to metazoan gap junctions (Weiss et al 2019).

Although a variety of cell-cell communication related proteins were identified and deeply investigated, their interplay in the process of assembling the communication apparatus is still poorly understood and the structural SJ components remain, with one exception, unknown (for reviews, see (Flores et al 2019, Herrero et al 2016, Kieninger & Maldener 2021)). This emphasizes the complexity of the cell-cell communication machinery and the importance of further investigations. Here, we describe the identification and mutational analysis of the protein SepN. By co-immunoprecipitation experiments using the plug-localized protein FraD as bait, SepN was identified as a potential FraD interacting protein. We show here, that SepN is itself involved in assembly of the SJs with particular importance in formation of the plug module. Furthermore, we suggest a critical role for the SJ plug in communication and SJ gating.

## Material and methods

### Cultivation of bacterial strains

All bacterial strains used in this study are described in suppl. Tab 1. Cyanobacterial strains were cultivated at 28 °C with constant illumination at 25-40 μE m^-2^ s^-1^ in liquid Bg11 medium (Rippka et al 1979) shaking at 100-120 rpm or on solidified Bg11 medium supplemented with 1.5% (w/v) Bacto agar. Liquid cultures of 700 mL for co-immunoprecipitation experiments were cultivated in bottles and bubbled with air, supplied with 2% CO_2_. Mutant strains grew in presence of the respective antibiotics at the following concentrations: 50 μg/mL neomycin, 5 μg/mL streptomycin and 5 μg/mL spectinomycin.

*Escherichia coli* (*E. coli*) strains were cultivated in LB medium at 37 °C supplemented with 50 μg/mL kanamycin, 25 μg/mL streptomycin and 100 μg/mL spectinomycin when indicated.

### Construction of mutant strains

DNA sequencing results were obtained from GATC biotech AG (Eurofins, Germany) and compared to reference sequences derived from the KEGG database (Kanehisa & Goto 2000). Purification of plasmid DNA and PCR fragments was performed using the ExtractMe Kit systems (Blirt, Poland). Triparental mating (conjugation) was performed to introduce plasmids into *Nostoc* cyanobacterial cells using the *E. coli* strains J53 (RP-4) (Wolk et al 1984) and HB101 (pRL528) carrying the cargo plasmid (Elhai and Wolk 1984). A description of used plasmids and sequences of oligonucleotides are summarized in supplementary table 1 and supplementary table 2, respectively.

As a control for α-GFP co-IPs, the gene encoding a version of the green fluorescent protein (GFP) *gfpmut2* (Cormack et al 1996) was cloned under the control of the *fraCDE* promoter region, yielding plasmid pIM800. P_*fraCDE*_*-gfpmut2* was amplified with oligonucleotides 1998/2235 using pIM779 (Weiss et al 2019) as template. The insert was ligated into EcoRI/BamHI digested shuttle vector pRL1049 (Black & Wolk 1994) and conjugated into *Nostoc*, creating strain 7120.800.

The *all4109* gene was interrupted by insertion of the cassette C.K3*t*4 provoking neomycin/kanamycin resistance under the control of the strong promoter P_*psbA*_ and a transcriptional terminator. For construction of this cassette, the earlier described cassette C.K3 (Elhai & Wolk 1988) was C-terminally fused downstream of the BamHI restriction site to the C-terminus of the bacteriophage T4 gene *32* (Krisch & Allet 1982), which harbors a translation tandem stop and a transcriptional terminator. The construct was ordered as a synthetic gene from Eurofins Genomics and amplified with oligonucleotides 1383/1384. To disrupt *all4109*, an upstream DNA fragment of *all4109* and a fragment within the end of the gene were amplified using oligonucleotides 2399/2397 and 2398/2400, respectively. The fragments were ligated via Gibson Assembly Cloning in a way flanking C.K3*t4* in XhoI/PstI digested pRL277 vector (Black et al 1995), yielding plasmid pIM825. After conjugal transfer of pIM825 into *Nostoc*, double-crossover homologous recombination with the genome and segregation of the resulting *all4109* mutant DR825 was checked via PCR using the oligonucleotides 2446/2447.

For localization studies, a C-terminal translational fusion of superfolder (sf) GFP to *all4109* was achieved by single homologous recombination of plasmid pIM834 with the genomic DNA. The coding sequence for sfGFP was amplified using oligonucleotides 1444/1445 and the *sfgfp-gene* bearing plasmid pIM660.2 as template (Bornikoel, unpublished). The C-terminal part of *all4109* was amplified without stop codon using oligonucleotides 2450/2451. Fragments were ligated via Gibson Assembly Cloning in XhoI/PstI digested vector pRL277 and the resulting plasmid pIM834 was transferred to *Nostoc* creating mutant SR834. Mutant segregation was checked via PCR with oligonucleotides 2446/2447, what only leads to amplification in the WT, since the whole plasmid is inserted into the tested region. To localize All4109 in a *fraD* mutant, single recombinant homologous recombination of pIM834 was performed in strain CSVT2 (Merino-Puerto et al 2010), creating the *all4109-sfGFP* expressing *fraD* mutant CSVT2.SR834.

To complement the *all4109* mutant, the predicted promoter region of the *all4110-all4109* operon (Softberry BPROM (Solovyev & Salamov 2011)) was amplified with oligonucleotides 2511/2512 and fused via PCR to the amplified *all4109* gene (oligonucleotides 2513/2514). The fusion product was ligated into EcoRI/BamHI digested pRL1049 (Black & Wolk 1994) via Gibson Assembly Cloning and the resulting plasmid pIM848 was conjugated into DR825, creating strain DR825.848.

### Co-immunoprecipitation and liquid chromatography mass spectrometry (LC-MS/MS)

Strains *Nostoc* WT, CSVT2 (Merino-Puerto et al 2010), CSVT2.779 (Weiss et al 2019), and 7120.800 were cultivated 7-10 days with supply of 2 % CO_2_ at 28 °C in constant light. For controls and specific conditions of independent immunoprecipitation experiments see supplement table 3. Cells were washed with PBS pH 7.4, resuspended in 15/20 mL PBS to OD_750_=15/25 (α-GFP-/α-FraD-pulldown) and crosslinked with 1 % glutaraldehyde for 30 min at RT when indicated. In one of the α-GFP immunoprecipitations crosslinking was performed with 0.6% formaldehyde for 30 min at RT followed by two washing steps and incubation for 10 min with 1 % glutaraldehyde at RT.

After washing with PBS and addition of a protease inhibitor cocktail tablet (cOmplete™, Roche), cells were lysed via three passages through a French press at 20,000 psi. Whole cells were discarded through low-speed centrifugation and membranes were collected via centrifugation at 48,000-70,000 g at 4 °C for 1 h. Membranes were solubilized in 500 μL 10 mM Tris-HCl, 100 mM NaCl pH 8 (α-GFP) or PBS pH 7.4 (α-FraD) supplemented with 1 % *N*-lauroylsarcosine for 1 h at RT and peptidoglycan was digested in parallel by addition of 1 mg/mL lysozyme. Insolubilized membranes were discarded after centrifugation at 21,000 g and 4 °C for 25 min. Co-IP with α-GFP magnetic beads (GFP-Trap®_MA, Chromotek) was performed following the instructions of the supplier. Solubilized membranes were incubated with the beads for 1:30 h under slow rotation at 4 °C. Bound proteins were eluted with 70 μL 2x SDS (sodium dodecyl sulfate) loading dye in Tris buffer (see above) and boiled at 95 °C for 10 min. For α-FraD immunoprecipitation, Dynabeads™ Protein G (Invitrogen) were incubated with 10 μg purified α-FraD antibodies in PBS for 10 min at RT. FraD antibodies were raised in rabbit against the synthetic peptide NH_2_-IWTGPTANPRGYFLRKSC-CONH_2_ within the periplasmic part of FraD (Pineda Antikörper-Service, Berlin, Germany). Solubilized membranes were incubated with α-FraD-bound magnetic beads for 1:30 h at 4 °C and eluted with 40 μL 50 mM glycine, pH 2.8 and 20 μL 6x SDS loading dye. Samples were boiled for 10 min at 70 °C and neutralized with 5 μL 1 M Tris, pH 9. Eluted proteins were separated on a 13 % SDS-PAGE gel. The gel was stained overnight with InstantBlue (Abcam), lanes were excised and in-gel digested with trypsin. LC-MS/MS analysis was performed using linear 60-min gradients and an Easy-nLC 1200 system coupled to a Q Exactive HF mass spectrometer (Thermo Fisher Scientific, Germany) by the Proteomcenter at the University of Tuebingen. Co-IPs with SR834 and DR825 as control were performed identically to α-GFP-FraD co-IPs.

Detected proteins with at least 100 times increased abundancy in the sample compared to the control were analyzed on their taxonomic distribution using STRING version 11 (Szklarczyk et al 2019) and HmmerWeb version 2.41.1. Proteins were considered as candidates for FraD-interacting proteins, if they showed a similar taxonomic distribution as FraD, which means conservation in filamentous cyanobacteria.

### *In silico* protein and gene analysis

Protein secondary structure prediction was performed using Protter webinterface version 1.0 (Omasits et al 2013). InterPro 82.0 (Mitchell et al 2018) and Pfam 33.1 (El-Gebali et al 2018) were used to scan for conserved domains.

### Light- and fluorescence microscopy

A Leica DM2500 B microscope with a Leica DFC420C camera was used for light microscopy. Fluorescence microscopy was performed with a 100x/1.3 oil objective lens of a Leica DM5500 B microscope connected to a Leica DFC360FX camera. GFP and chlorophyll fluorescence were excited using a BP470/40 nm or BP535/50 nm filter and emission was monitored using a BP525/50 nm or BP610/75 nm filter, respectively. Z-stacks with 0.1 μm intervals were taken to perform three-dimensional deconvolution using the built-in function of the Leica ASF software.

For quantification of the fluorescence in the septum, each xy pixel of 20 slices of a raw (not deconvoluted) z-stack was summed into one image and the mean intensity of septal ROIs was measured. Background fluorescence was measured with identical ROIs in the cytoplasm. The averaged background fluorescence was subtracted from every single septum measurement and used for normalization of the values.

### FRAP and CCCP-FRAP

FRAP and CCCP-FRAP experiments were performed as described previously (Weiss et al 2019). In short, 500 μL cells were stained with 8 μL of a calcein acetoxymethylester solution (1 mg/mL in DMSO) and incubated for 90 min the dark at 28 °C. After three washing steps with Bg11 medium, cells were incubated either with (50 μM CCCP) or without protonophore for further 90 min in the dark. 10 μL of the stained cells were spotted onto a Bg11 agar plate for imaging with excitation at 488 nm of a laser at 0.2 % intensity of a Zeiss LSM 800 confocal microscope using a 63x/1.4 oil-immersion objective and the ZEN 2.6 (blue edition) software. Calcein fluorescence emission was detected at 400-530 nm simultaneously with chlorophyll autofluorescence emission at 650-700 nm. Bleaching of a specific cell was achieved by shortly increasing the laser intensity to 3.5 % after 5 pre-bleached images were taken. Recovery of fluorescence in the bleached cell was monitored in 1 s intervals for 30-180 s. Analysis of the acquired images was done with ImageJ (version 1.51j) and GraphPad Prism 6 as described earlier (Weiss et al 2019).

### Isolation of septal peptidoglycan and transmission electron microscopy

Septal PG was isolated after Kühner et al (Kühner et al 2014) with some changes. Cells grown for 3 d on Bg11 agar plates were resuspended in 700 μL 0.1 M Tris-HCl pH 6.8 and sonicated (Branson Sonifier 250) for 2 min at a duty cycle of 50 % and output control 1. After addition of 300 μL 10 % SDS, the samples were boiled for 30 min at 99 °C with shaking at 300 rpm. The suspension was washed two times with ddH_2_O (5 min, 10,000 rpm), incubated for 30 min in a sonifier waterbath. After washing with ddH_2_O, the debris was resuspended in 1 mL 50 mM Na_3_PO_4_ pH 6.8 and incubated 3 h at 37 °C with 300 μg α-Chymotrypsin under constant rotation. After that, α-Chymotrypsin was added again and the samples were incubated over night at 37 °C. The next day, the enzyme was heat inactivated for 3 min at 99 °C. After another sonication (30-90 s) and washing step with ddH_2_O, the PG septa were resuspended in 50-500 μL ddH_2_O. All solutions were filtered with a 0.22 μm filter.

10 μL of isolated septal PG suspension was incubated on an UV-irradiated (16 h) formvar/carbon film-coated copper grid (Science Services GmbH, Munich, Germany) for 10-30 min. Samples were stained with 1 % (w/v) uranyl acetate and imaged with a Philips Tecnai10 electron microscope at 80 kV and a Rio Camera (Gatan).

### Plunge freezing of *Nostoc* cells

*Nostoc* cultures were concentrated by gentle centrifugation (1000 x g, 5 min) and removing 2/3 of the medium. 3.5 μL of cell suspension was applied on glow-discharged copper or gold EM grids (R2/2, Quantifoil) and automatically back-blotted (Weiss et al 2017) for 4-6 s, at 22ºC with 100% humidity and plunged into liquid ethane/propane (Tivol et al 2008) using a Vitrobot plunge freezing robot (ThermoFisher Scientific, Waltham, MA) Frozen grids were stored in liquid nitrogen.

### Cryo-Focused Ion Beam milling

Thin lamellae through plunge frozen *Nostoc* filaments were obtained by automated sequential FIB milling according to (Zachs et al 2020). Briefly, EM grids were clipped into FIB milling autoloader-grids (ThermoFisher Scientific, Waltham, MA) and mounted onto a 40º pre-tilted grid holder (Medeiros et al 2018) (Leica Microsystems GmbH, Vienna, Austria) using a VCM loading station (Leica Microsystems GmbH, Vienna, Austria). For all grid transfers under cryo-conditions, a VCT500 cryo-transfer system (Leica Microsystems GmbH, Vienna, Austria) was used. Grids were sputter-coated with a ∼4 nm thick layer of tungsten using ACE600 cryo-sputter coater (Leica Microsystems GmbH, Vienna, Austria) and afterwards transferred into a Crossbeam 550 FIB-SEM dual-beam instrument (Carl Zeiss Microscopy, Oberkochen) equipped with a copper-band cooled mechanical cryo-stage (Leica Microsystems GmbH, Vienna, Austria). The gas injection system (GIS) was used to deposit an organometallic platinum precursor layer onto each grid. Grid quality assessment and targeting of the cells were done by scanning EM (SEM) imaging (3-5 kV, 58 pA). Coordinates of chosen targets were saved in the stage navigator and milling patterns were placed onto targets’ FIB image (30 kV, 20 pA) using SmartFIB software. To mill 11 μm wide and 200 nm thick lamellae a total of four currents were used and currents were gradually reduced according to lamella thickness [rough milling (700 pA, 300 pA, and 100 pA) and polishing (50 pA)]. After the milling session, the holder was brought back to the loading station, grids were unloaded and stored in liquid nitrogen.

### Cryo-electron tomography

Lamellae through *Nostoc* filaments were examined by cryo-electron tomography. Data were collected on a Titan Krios 300kV FEG transmission electron microscope (ThermoFisher) using a Quantum LS imaging filter (slit width 20 eV) and K2 Summit direct electron detector (Gatan). A low magnification (135x) overview of the grid was recorded using SerialEM (Mastronarde 2005) to identify lamellae. Tilt series were collected automatically using a custom-made SerialEM script. Target tracking was achieved via cross-correlation to the previous record image. For focusing, a single autofocus routine was performed at zero degree and focus change over the entire tilt series was estimated using the focus equation from UCSF tomography (Zheng et al 2007).

Tilt series covered an angular range from −50° to +70° at 2° increments with a defocus of −8 μm, total accumulated dose of ∼120 - 140 e^-^ / Å^2^ and a pixel size of 3.45 Å. Tomogram reconstruction was performed according to (Weiss et al 2017) using IMOD package (Mastronarde & Held 2017). A deconvolution filter (Tegunov & Cramer 2019) was used for tomograms shown in figures.

### Subtomogram averaging

Subtomogram averaging was performed using Dynamo (Castaño-Díez et al 2012). Briefly, SJs were identified visually in individual tomograms and their coordinates were saved as “oriented particles” models in the catalogue. For the initial average, 418, 282, and 89 particles were picked for WT (open state), WT (closed state), and *sepN* mutant respectively. To exclude reference bias, first templates were created by averaging fifty randomly picked particles for each dataset. Particles were aligned for three iterations using 2×2-binned tomograms [box-size 72 × 72 × 72 pixels, alignment mask: sphere (radius = 20 pixels)]. To attenuate a prominent missing wedge in the average due to preferential particle orientation, we randomized the rotational angle using “dynamo_table_randomize_azimuth(table)” matlab command. Another round of 4 iterations was performed and obtained averages and tables were used for refinement. For the final refinement run (9 iterations with a tight alignment mask), we applied five-fold symmetry according to results in (Weiss et al 2019). The final, 5-fold symmetrized averages resulted from 387 (WT open state), 248 (WT closed) and 87 (*sepN* mutant). Estimation of the resolution for each dataset was generated by using sg calculate_FSC script from (Chen et al 2013). The difference map between subtomogram averages of wild type closed state and *sepN* mutant was generated with the diffmap program (https://grigoriefflab.umassmed.edu/diffmap). 3D rendering and coloring of different SJ modules were done with Chimera. Example tomograms and subtomogram averages have been uploaded to the electron microscopy databank (Accession codes: EMD-XXXX – EMD-XXXX)

### Immunolocalization of FraD

Immunolocalization of FraD in *Nostoc* strains was principally performed as described by Büttner et al. with some minor changes (Büttner et al 2016). *Nostoc* filaments were grown for three days on Bg11 plates and resuspended in 1 mL PBS to OD_750_=1. After three washing steps, the cells were fixed with 1 mL of HistoChoice Tissue Fixative (Sigma-Aldrich) for 10 min at RT and 30 min at 4 °C. Cells were washed three times with PBS, once with 70 % EtOH prechilled at -20 °C and again with PBS. Next, the cells were treated with 1 mg/mL lysozyme in 1 mL GTE buffer (50 mM glucose, 20 mM Tris-HCl pH 7.5, 10 mM EDTA) for 5 min at RT, resuspended in 200 μL PBS and dropped onto a Polysine slide (ThermoFisher Scientific, Germany). Cells were dried at RT and rehydrated with 200 μL PBS for 5 min. After blocking with 200 μL 2 % w/v BSA in PBS for 20 min, cells were incubated overnight in a wet chamber at 4 °C with 10 μg/mL α-FraD antibodies (see above) in BSA-PBS. Cells were washed five times with PBS and incubated 2 h at RT in the dark with FITC-coupled α-rabbit antibodies (1:200 in BSA-PBS, Sigma Aldrich). After washing and drying, one drop of Vectashield Mounting Medium H-1200 (Vector Laboratories, USA) was applied, covered with a coverslip and sealed with nail polish. Fluorescence was imaged using a DM5500B Leica microscope and a DFC360FX monochrome camera. Autofluorescence was detected as described before, FITC-fluorescence was detected by using a BP470/40 nm excitation filter and a BP525/50 nm emission filter. Z-stacks with 0.1 μm intervals were taken and applied to the Leica ASF built-in function for 3D-deconvolution.

### Test on significance

Statistical analysis was performed with GraphPad Prism version 6.01. Statistical significance of two groups was tested using an unpaired Student’s *t*-test. Comparison of one group with multiple other groups was performed via ordinary one-way ANOVA followed by Dunnett’s multiple comparison test. Significance *P* is indicated with asterisks: *: *P*≤0.05; **: *P*≤0.01; ***: *P*≤0.001; ****: *P*≤0.0001; ns (not significant): *P*>0.05.

## Results

### Identification of SepN as putative interaction partner of FraD

Recently, we verified FraD as the first known structural SJ component with localization to the plug module (Weiss et al 2019). In order to identify further proteins of the multimeric SJ protein complexes, we performed co-immunoprecipitations (co-IPs) using FraD or GFP-FraD as bait. All independently performed co-IPs (in total 6) are summarized in Table S3. Detected proteins were considered as putatively interacting with FraD, when they were 100 times more abundant in the sample than in the control and if they were conserved only in filamentous cyanobacteria similarly to FraD. The most promising candidate was the protein encoded by gene *all4109*, since it was detected in all crosslinked and non-crosslinked samples in every co-IP experiment with high abundancy. Furthermore, the protein was described as a signature protein for the order Nostocales (Gupta & Mathews 2010). We renamed All4109 to SepN and will use the latter name hereafter.

The gene *sepN* (*all4109*) is the second in a predicted operon downstream of *all4110* (MicrobesOnline Operon Prediction). Whereas the gene product of the latter is annotated as magnesium transport protein CorA that mediates the influx of magnesium ions, *all4109* encodes an unknown protein of 235 aa. However, an internal promoter sequence in the upstream gene *all4110* was observed previously and could allow single transcription of *all4109* (Mitschke et al 2011). All4109 (SepN) is predicted to contain one transmembrane domain (aa 7-29, InterPro), with the N-terminal end facing the cytoplasm and the C-terminal part ranging into the periplasm (Protter, see methods). Pfam domain analysis revealed poor similarity of the N-terminal part to an Apo-citrate lyase phosphoribosyl-dephospho-CoA transferase (E-value 0.0049). Gene co-occurrence with FraD is stated by the STRING database, which also reports conserved homologs of *sepN* in filamentous cyanobacteria of the *Oscillatoriaphycideae, Nostocaceaae*, and *Stigonematales*. The predicted absence of enzymatic domains and the FraD-like taxonomic distribution in filamentous cyanobacteria fit to an expected role in cell-cell communication.

### SepN localizes to the septum of adjacent cells

Proteins related to the cell-cell communication system of *Nostoc*, like FraC, FraD, SepJ, AmiC1 and SjcF1, localize in the septum between neighboring cells (Berendt et al 2012, Flores et al 2007, Merino-Puerto et al 2010, Merino-Puerto et al 2011, Rudolf et al 2015). To visualize the subcellular localization of SepN, a single recombinant C-terminal fusion to superfolder (sf) version of GFP was inserted into the genome replacing the wild-type gene. In this created strain SR834, the GFP-fusion protein was expressed from its native promoter avoiding overexpression artefacts. A superfolder version of GFP was used to ensure correct folding of GFP in the periplasm. Analysis of the *sepN-sfgfp* strain via fluorescence microscopy revealed focused fluorescence foci in the septa of adjacent vegetative cells similarly to GFPmut2-FraD (CSVT2.779) (Fig. 1A). As fluorescence background control, WT cells not expressing any GFP were imaged with identical settings. SepN was absent in constricting septa of dividing cells but appeared in all mature septa (Fig. 1B).

**Figure 1:**
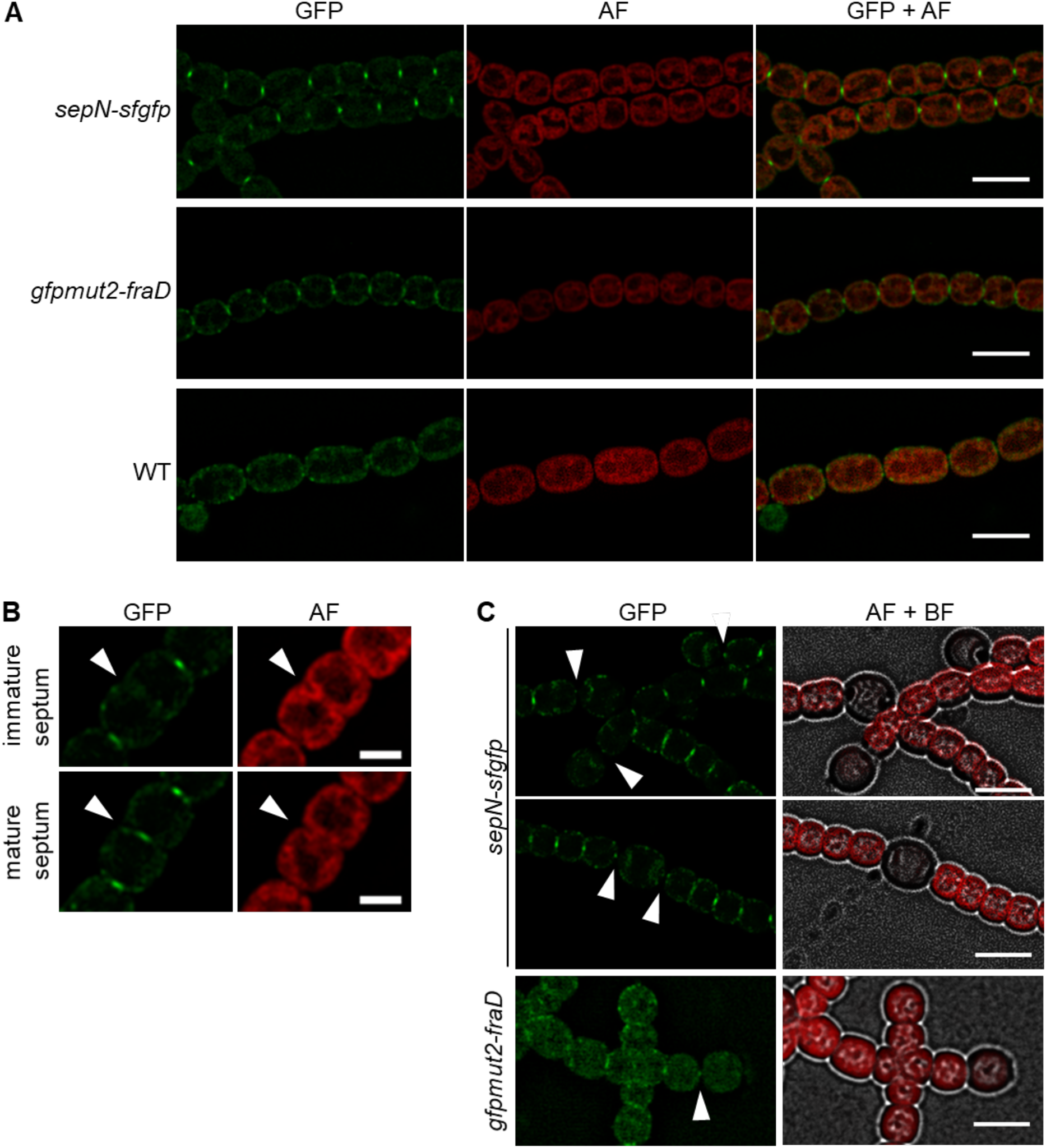
Subcellular localization of SepN-sfGFP. (A) Green fluorescence foci of SepN-sfGFP (SR834) were detected in the septum between vegetative cells similarly to GFPmut2-FraD (CSVT2.779). WT cells were included as a negative control. Shown are fluorescence micrographs of GFP, autofluorescence (AF) coming from the photosynthetic pigments and an overlay of both. Scale bar, 5 μm. (B) No fluorescence signal could be detected for SepN-sfGFP in immature septa of dividing cells, but appeared early after cell division. Arrow heads point to the septum of a dividing cell. Bar, 2 μm. (C) SepN-sfGFP was detected in the septum of terminal (upper panel) and intercalary (middle panel) heterocysts. The *gfpmut2-fraD* strain CSVT2.779 was included for comparison. Cultures were grown for 4 d on nitrogen-depleted agar plates before imaging. Scale bar, 5 μm. Arrow heads point to the heterocyst-vegetative cell septa. AF, autofluorescence; BF, bright field.

In mature heterocysts of filaments, which had been cultured for four days after nitrogen stepdown one small fluorescent spot was detected in the heterocyst-vegetative cell septum of both, terminal and intercalary heterocysts (Fig. 1C). A dispersed fluorescence signal for SepN-sfGFP was observed surrounding the polar plugs (also known as nodules), which are intracellular assemblies of the nitrogen storage molecule cyanophycin (multi-L-arginyl-poly(L-aspartic acid). They accumulate at the poles of the heterocysts (Jäger et al 1997).

### Cell-cell communication and gating of septal junctions was impaired in a *sepN* mutant

Several mutants lacking proteins involved in cell-cell communication exhibit a lower rate of molecular exchange compared to the WT, due to a reduced number of nanopores and SJs (Bornikoel et al 2017, Bornikoel et al 2018, Kieninger & Maldener 2021, Merino-Puerto et al 2010, Nürnberg et al 2015). To get a first hint on potential involvement of SepN in intercellular communication, we traced the cell-to-cell diffusion of the fluorescent dye calcein by FRAP measurements in *sepN* and *sepN-sfGFP* mutants. We constructed a disruption mutant (DR825) by insertion of the neomycin resistance cassette C.K3t4 into the *sepN* gene and introduced the *sepN-sf*GFP fusion by single homologous recombination with the genome (SR834) (see methods section).

The FRAP measurements showed that the fluorescence recovery rate constant *R* in the *sepN* mutant was significantly reduced, which is also true for the *fraD* mutant (Fig. 2A). Complementation of the *sepN* mutant with SepN expressed from a self-replicating plasmid under control of its native promoter (strain DR825.848) restored a WT-like communication rate. Interestingly, the *sepN-sfgfp* strain showed a similar *R* as the knockout mutant. Slower dye exchange of the GFP-fusion strain might be a hint on steric hinderance of diffusion or SepN interaction partners by GFP, similarly to the recently described GFP-FraD mutant (Weiss et al 2019).

**Figure 2:**
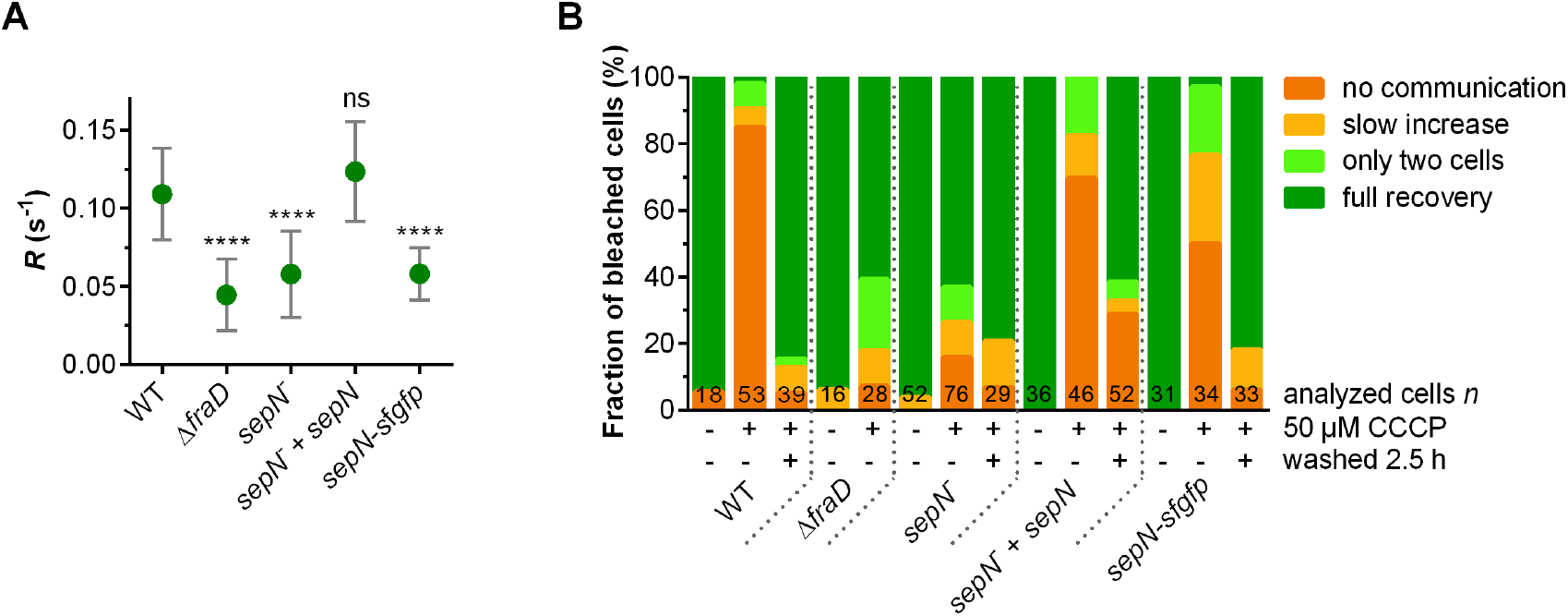
CCCP-FRAP analysis of the sepN mutant.(A) The fluorescence recovery rate constant R in FRAP experiments was calculated for full recovering, untreated cells. Data for CSVT2 was taken from (Weiss et al., 2019). (B) FRAP-responses of calcein-stained untreated, CCCP-treated (90 min, 50 μM), and washed cells after CCCP treatment were assigned to one of four groups indicated by the color scheme. The numbers within the bars indicate the number of analyzed cells (n) from different filaments. Cumulated results from at least two independent cultures are shown. Results of two independent sepN mutant and complemented strains were cumulated. Data for CSVT2 were taken from (Weiss et al 2019). Strains: ΔfraD, CSVT2; sepN-, DR825; sepN-+ sepN, DR825.848; sepN-sfgfp, SR834.

As we showed previously, SJs are gated cell-cell connections and closure can be triggered by disruption of the proton motive force (Weiss et al 2019). In this setup, cells were treated with the protonophore CCCP prior to FRAP analysis. We furthermore showed, that a *fraD* mutant was insensitive to the CCCP treatment, which manifested in a major fraction of still communicating cells (Weiss et al 2019). Inability to regulate intercellular communication was linked to the absence of the SJ cap and plug modules in the *fraD* mutant. In the WT, the cap undergoes a conformational change upon stress (Weiss et al 2019). To investigate the impact of SepN on SJ gating, we performed CCCP-FRAP experiments with the *sepN* mutant, the complemented mutant and with the GFP-fusion strain. The FRAP responses were assigned to one of four groups as described previously, which are “no communication”, “slow increase” of fluorescence recovery”, recovery only between “two cells” and “full recovery” (Weiss et al 2019). Communication under standard growth conditions was possible in all mutants, since the majority of calcein-stained cells showed a full recovery response after bleaching (Fig. 2B). Treatment of the *sepN* mutant with 50 μM CCCP led to a “no communication” response in only 16 % of the analyzed cells, which was in contrast to 85 % closing in the WT. A “full-recovery” phenotype in 63 % of the CCCP treated cells of the *sepN* mutant was very similar to the Δ*fraD* strain, which showed a fraction of 61 % communicating cells after CCCP treatment. Accordingly, gating of SJs was strongly affected in the absence of SepN suggesting a role of this protein as assembly factor or even structural element of the SJ complexes. Since SJ closure was restored in the complemented mutant, the phenotype was indeed caused by lack of the SepN protein. However, reopening of SJs after removal of CCCP was not as efficient as in the WT. SR834, in which *sepN* was exchanged for *sepN-sfGFP*, was able to gate its SJs, however, the fraction of non-communicating cells upon CCCP treatment was by 35 % reduced in comparison to the WT (Fig. 2B). The fusion to GFP might therefore interfere with other structural components of the SJs like FraD or sterically affect the process of SJ closure.

### The nanopore array of a *sepN* mutant is similar to the WT

To further investigate the role of SepN during SJ assembly, we set out analyze the nanopore array of *sepN* mutant filaments. The nanopore array is an essential feature of the cell-cell communication system, since it builds the scaffold for the SJ protein complexes. As shown earlier for several mutants, the rate of molecular diffusion depends on the amount of nanopores per septum (reviewed in (Kieninger & Maldener 2021)). Since the diffusion rates in the *fraD* mutant (as already described in (Merino-Puerto et al 2011)) and the *sepN* mutants were severely reduced (Fig. 2B), we purified septal PG discs and compared the respective nanopore array to WT septa. As expected, deletion of *fraD* led to a greatly diminished number of nanopores per septum (Fig. 3A, B). Surprisingly, disruption of *sepN* or fusion to GFP did not lead to an altered nanopore array compared to the WT. The diameter of the isolated PG disks was significantly enlarged in a *fraD* mutant, but similar to the WT in the absence of SepN (Fig. 3C). Whereas the nanopores in septa of the *fraD* mutant were strongly enlarged compared to the WT, their diameter was only slightly bigger in a *sepN* mutant (Fig. 3D).

**Figure 2:**
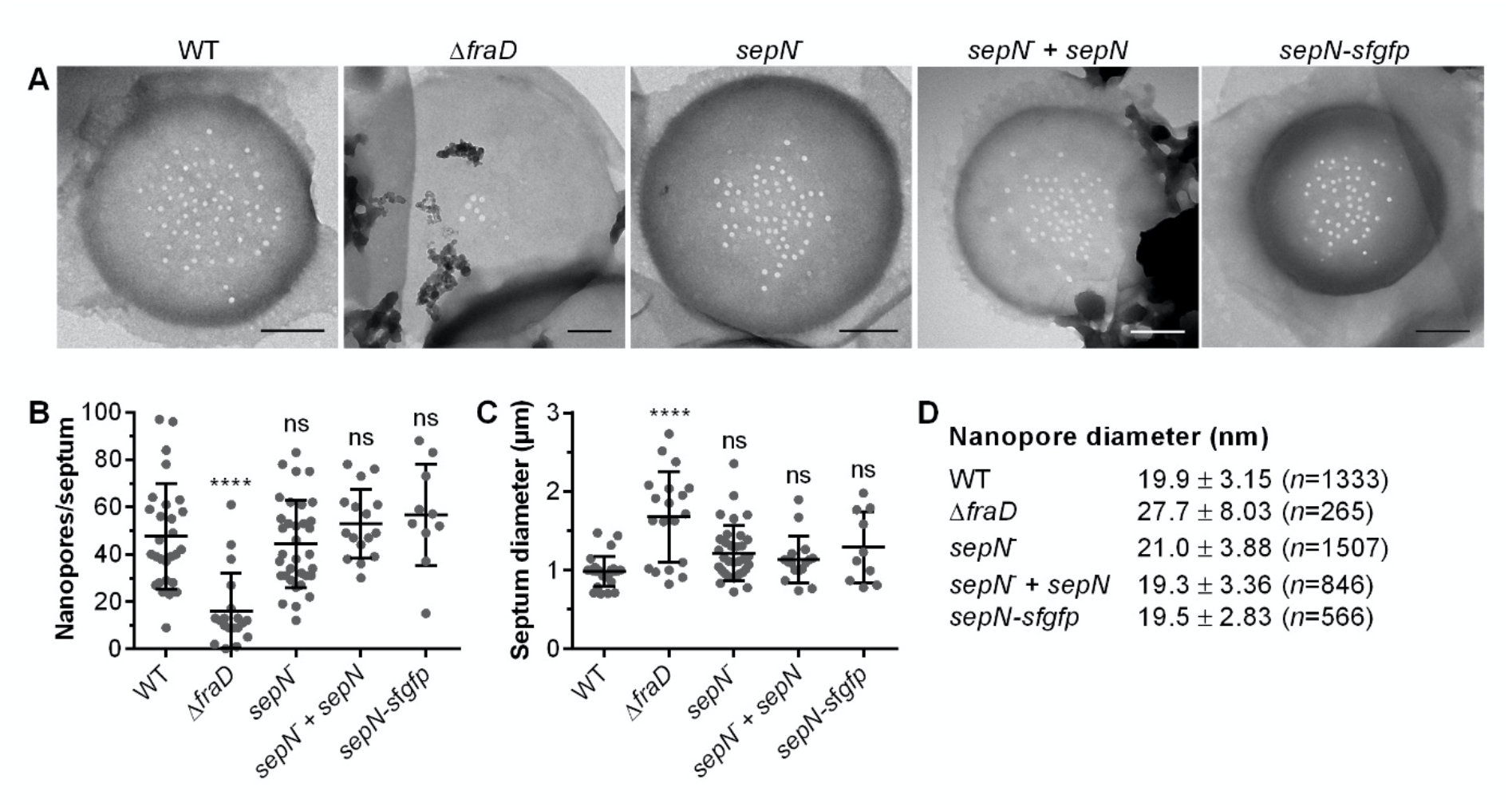
Disruption of *sepN* does not reduce the nanopore array.Septal PG discs were isolated and analyzed via transmission electron microscopy (A) Representative transmission electron micrographs of the indicated strains. Scale bar, 250 nm. (B) The mean and standard deviation of nanopores per septum and (C) the septum diameter was measured. Each dot represents an analyzed PG septum. (D) The mean and standard deviation of the nanopore diameter cumulated from all analyzed septa is shown. The number of analyzed nanopores *n* is indicated. Data for the *sepN* mutant were cumulated from two independent mutant clones. Strains: Δ*fraD*, CSVT2; *sepN*^-^, DR825; *sepN*^-^ + *sepN*, DR825.848; *sepN sfgfp*, SR834.

### *sepN* mutant is lacking the plug module

Since the reduced communication rate in the *sepN* mutant seemed not to be a result of fewer nanopores, we assumed that SepN might be a structural component of SJs. As cryo-electron tomography (cryoET) already demonstrated to be a suitable tool to study structural changes of SJs *in situ* (Weiss et al 2019), we next employed cryoET to investigate whether the phenotype observed in the *sepN* mutant is caused by alterations in SJs’ architecture. Plunge frozen *sepN* mutant filaments were thinned by cryo-focused ion beam (FIB) milling (Marko et al 2007, Villa et al 2013), and tomograms of individual septal areas were acquired on the resulting lamellae (*n* = 31 tomograms). Analysis of individual tomograms as well as the direct comparison with WT SJs revealed striking differences in the cap/plug area, whereas the most pronounced difference is a missing density for the plug (Fig. 4A). For a detailed analysis of the differences in SJ architecture of the *sepN* mutant, we applied subtomogram averaging on 98 SJ ends. The resulting average revealed that SJs in the *sepN* mutant are indeed missing a plug density (Fig 4B, middle). The shape of the cap resembled the previously described closed state of WT SJs (Weiss et al 2019), which is characterized by a narrower cap module compared to the WT open state as well as the disappearance of openings between the cap’s arches (Fig. 4B). A difference map calculation (Fig.4C) between SJs of the *sepN* mutant and WT in a closed state further emphasized the difference in the plug area and highlighted the similar cap architecture without any detectable openings (Fig. 4D). This observation, together with the inability of gating communication upon CCCP treatment, indicates that the cap alone cannot seal the SJs completely and the presence of the plug is required for proper closure of septal junctions.

**Figure 4:**
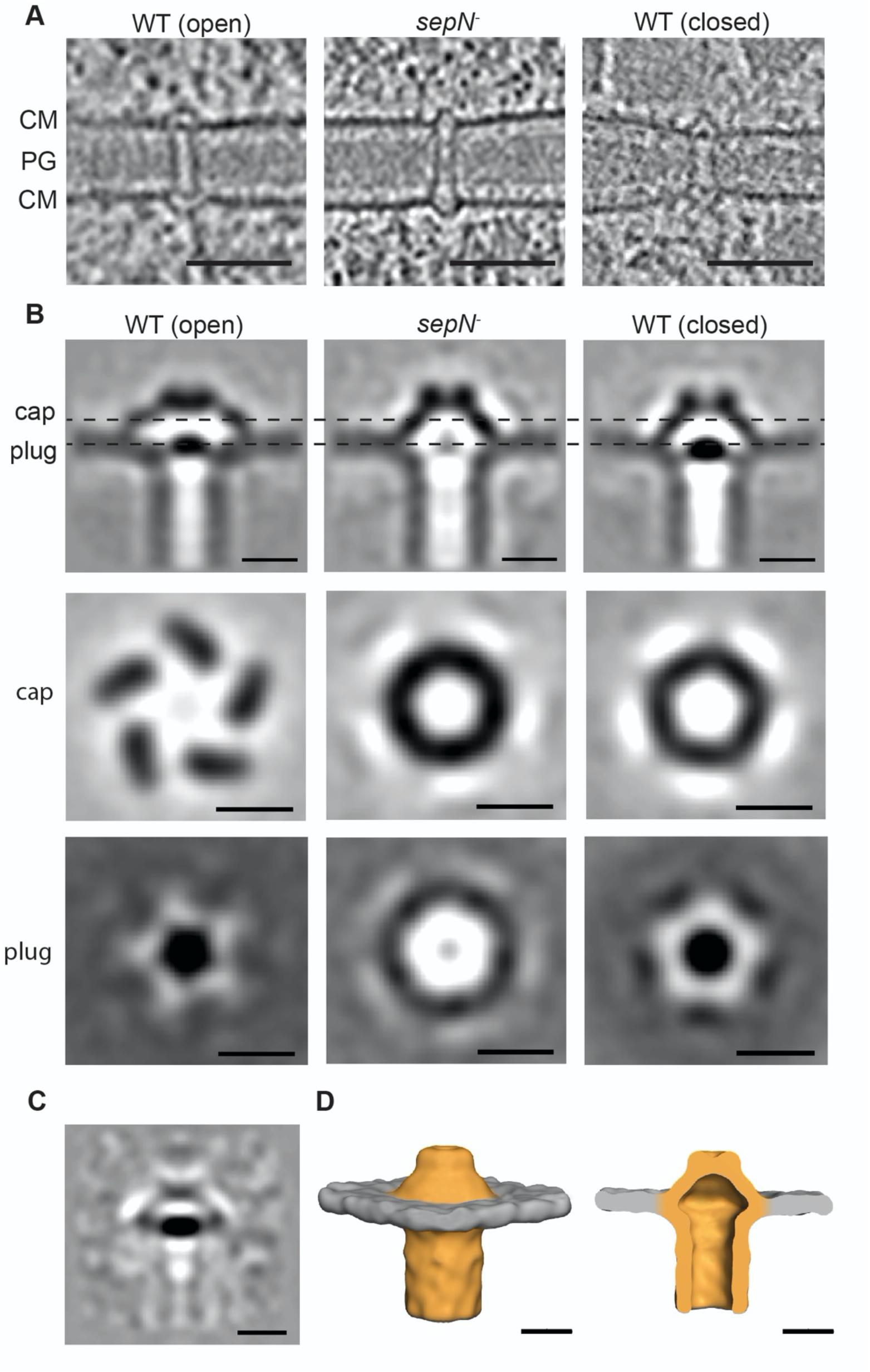
SJs from the *sepN* mutant show major differences in the architecture of the cap/plug module compared to the WT (A) Shown are 13.5 nm-thick slices through cryotomograms showing septum-spanning SJs in wild-type (untreated, open SJ state, left), *sepN* mutant (middle) and wild-type cells after CCCP treatment (closed SJ state, right). CM, cytoplasmic membrane; PG, septal peptidoglycan. Bar, 50 nm. (B) Subtomogram averaging of the *sepN* mutant SJs (middle) revealed a missing plug module. The cap of the *sepN* mutant SJs was resembling the closed state of wild-type SJs (right; for comparison is the open state is shown on the left). Shown are longitudinal and cross-sectional slices (0.68 nm) through the averages. Sliced positions are indicated by dashed lines. Bars, 10 nm. (C) A difference map calculation between the subtomogram averages of SJs from CCCP-treated wild-type cells (closed SJ state) and *sepN* mutant cells revealed that the most prominent difference is the missing density in the plug region. Bar, 10 nm. (D) Surface representation of the subtomogram average of the *sepN* mutant SJs showed no openings in the cap module, resembling the shape of WT SJs in the closed state but without plug module. Bars, 10 nm

### FraD localizes properly in a *sepN* mutant

To obtain further insights in the connection between FraD and SepN, the subcellular localization of FraD was investigated in a *sepN* mutant and vice versa. For this, we first visualized the septal localization of FraD in the WT and in the *sepN* mutant background via immunolocalization, using α-FraD and a FITC-coupled α-rabbit secondary antibody, showing a similar distribution of GFP in both strains (Fig. 5A). In some septa of the *sepN* mutant somewhat broader distribution compared to the WT was observed. Next, we aimed to localize SepN in a *fraD* mutant, which is why the plasmid pIM834, which codes *sepN-sfgfp*, was single recombinantly inserted into the Δ*fraD* strain yielding strain CSVT2.SR834. The latter was analyzed via fluorescence microscopy with the exact same settings as the *sepN-sfgfp* fusion strain in the WT background (Fig. 5B). Interestingly, the focused fluorescence foci in the septum observed for SepN-sfGFP in the WT background were absent in the majority of Δ*fraD* cells. Consequently, localization of SepN might be dependent on the presence of FraD, but not vice versa.

**Figure 5:**
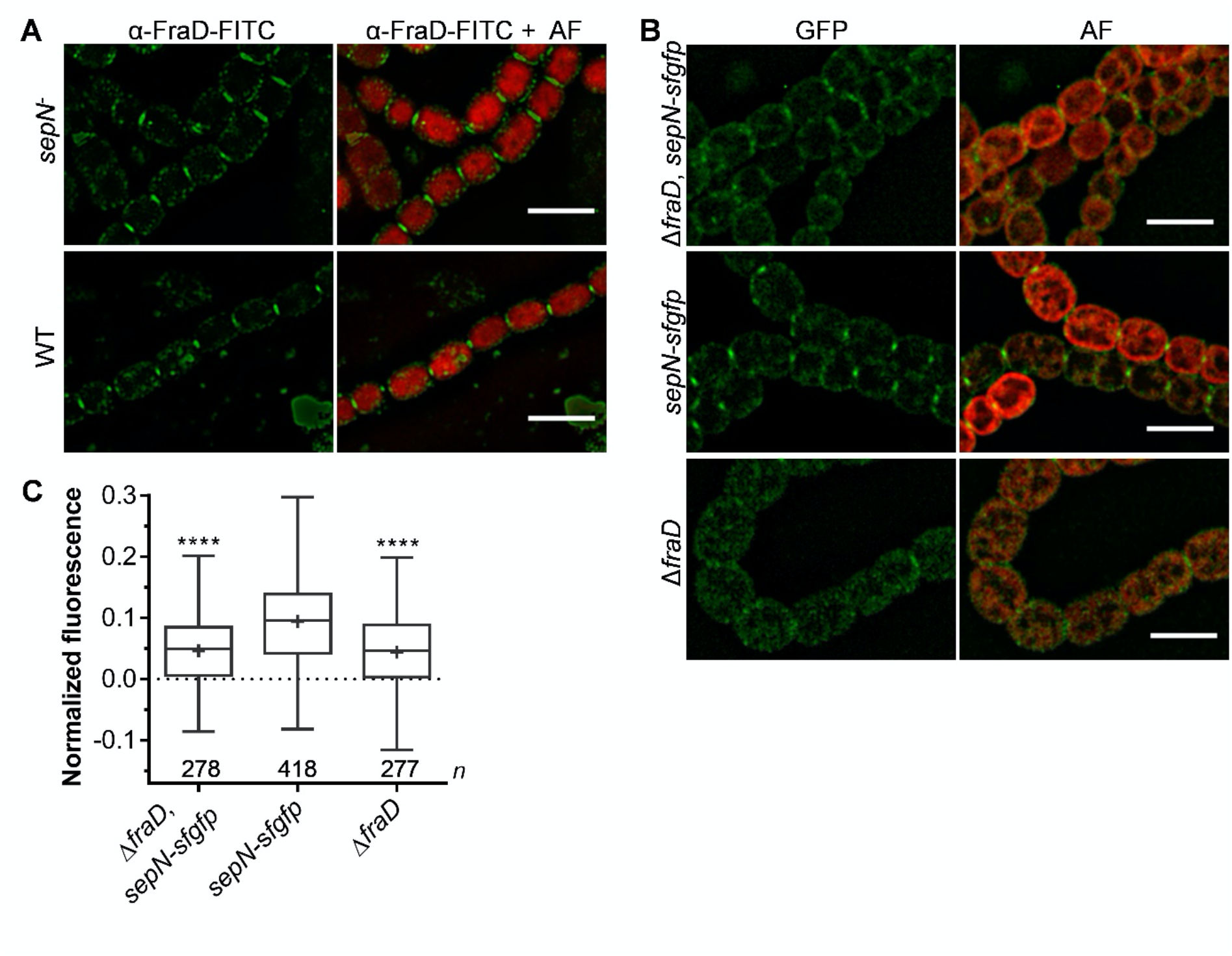
Mutual influence of SepN and FraD on their localization. (A) Immunolocalization of FraD in the *sepN* mutant DR825 and the WT was performed using primary α-FraD and FITC-coupled secondary antibodies. Representative 3D-deconvoluted fluorescence images are shown. Scale bar, 5 μm. FITC, fluorescein isothiocyanate; AF, autofluorescence. (B) Plasmid pIM834 encoding for *sepN-sfgfp* was single recombinantly inserted into the genomic *sepN* gene of the *fraD* mutant (CSVT2) or WT background. Emission of GFP was analyzed via fluorescence microscopy. Representative 3D-deconvoluted micrographs are shown. Contrast was enhanced for better visualization. Scale bar, 5 μm. (C) The mean fluorescence in the septa of the GFP-fusion strains shown in (B) was quantified as described in the methods. Box and whiskers projections show min to max values, the line in the box marks the median. The numbers of measured septa (*n*) are indicated below the boxes.

### FraD is the most abundant protein in a co-IP with SepN-sfGFP

Immunoprecipitation of membrane fractions using SepN-sfGFP as bait was performed with the intention to detect FraD as further evidence for their interaction. Immunoprecipitation was performed with α-GFP antibodies as described for GFP-FraD. A glutaraldehyde crosslinked and non-crosslinked sample, respectively, was used for co-IP and the *sepN* knockout mutant served as negative control. Indeed, in both, crosslinked and non-crosslinked sample, FraD was the most abundant protein beside the bait SepN itself. This gives further evidence, that SepN and FraD are interacting proteins. Interestingly, FraC was detected in both, the cross-linked and the non-crosslinked sample among the twenty most abundant proteins. Despite FraC is encoded in an operon with FraD, it was never detected in co-IPs directed against the latter.

## Discussion

In the past decade, several proteins were investigated that are involved in formation of the nanopore array and SJ complexes to set up a communication system in multicellular cyanobacteria (for reviews see (Flores et al 2019, Kieninger & Maldener 2021). However, until now, only FraD could be assigned as component of the SJs with localization to the plug module, which emphasizes the complexity of the establishment of the intercellular communication machinery. In order to gain a deeper understanding of these multimeric protein complexes, we aimed to identify further SJ components. Since SJs show functional but not structural analogy to metazoan gap junctions, SJ forming proteins cannot be found based on genomic blast analysis. In this study, we used FraD as bait to co-immunoprecipitate interacting proteins. Here, we focused on investigation of the most prominent hit detected in all conducted co-IPs, SepN.

A fluorescence-tagged version of SepN allowed us to visualize its septal localization as a focused spot between vegetative cells (Fig. 1). This characteristic localization made us name All4109 SepN referring to septal protein SepJ (Flores et al. 2007). Septal localization of SepN-sfGFP was also found between heterocyst and vegetative cells, however in lower intensity. This might be due to the restricted heterocyst septum and therefore less SJ proteins in the septal plane. The SepN-sfGPF signal could only be detected in mature septa, but not at sites of Z-ring formation, where cell division starts. The nanopore array is not drilled until the septum is fully closed, which may explain also the absence of the SJ proteins in immature septa, which have not yet completed the nanopore array. This is also true for the SJ protein FraD, which is absent in constricting septa (Merino-Puerto et al 2010). In contrast, the septal proteins, which are not structural parts of SJs like AmiC (Berendt et al 2012), SepJ (Flores et al 2007), FraC (Merino-Puerto et al 2010) and SepI (Springstein et al., 2020), migrate with the FtsZ ring during septum constriction and may have a function in maturation of the nanopore array (Kieninger & Maldener 2021).

Recently, we showed that intercellular communication was blocked upon stress, and demonstrated a reversible conformational change of the cap module (Weiss et al 2019). Failure of SJ gating is a strong hint that the SJ protein complex might be incompletely or aberrantly formed, as it was the case for the *fraD* mutant. In this mutant, SJ cap and plug modules were absent, suggesting a role of these modules in SJ closure. Disruption of *sepN* led to a strongly decreased ability of SJ closure, similarly to the Δ*fraD* strain (Fig. 2A). We therefore concluded that SepN is essential for SJ assembly and might be a structural part of these complexes. Despite the *sepN* mutant exhibited a reduced rate of molecular exchange compared to the WT (Fig. 2B), it harbored a WT-like nanopore array except for slightly wider nanopores (Fig. 3). The combination of reduced communication rate and a WT-like nanopore array was an additional hint, that intercellular communication might be slowed down due to structurally reasons of the communication channels. This assumption was confirmed by cryo-electron tomograms of the *sepN* mutant, in which the plug module was absent. Interestingly, and in contrast to the *fraD* mutant, the cap module was present. However, it rather resembled the closed cap state in the WT, because lateral openings were absent. We hypothesize, that the reduced rate of intercellular communication is caused by this closed-like cap. Since full inhibition of molecule diffusion was not possible in this mutant, the plug plays a key role in sealing SJs and interrupting cell-cell communication. Furthermore, the plug might also be important for keeping the cap structure in that particular conformation that was described as “open state” (Weiss et al. 2019). However, we cannot exclude that SepN interacts with cap components and controls the cap conformation, which is needed for plug formation.

Interestingly, fusion of sfGFP to SepN reduced the rate of intercellular communication to a similar level as in the knockout strain suggesting that the large GFP-tag interfered with a SJ-module conformation that allows molecule transfer. Cryotomograms and subtomogram averages, however, did not reveal an obvious difference in SJ architecture compared to wild type SJs. The size of sfGFP is at the resolution limit of an individual tomogram and since the tag might show a high flexibility, also the absence of an additional density in the subtomogram average does not exclude a direct involvement of SepN in the SJ complex. Previously, an extra density of GFP was observed in a SJ subtomogram average of a GFP-FraD fusion strain, because the fusion protein was localized in the SJ tube lumen (Weiss et al 2019).

Strikingly, septal localization of FraD in the *sepN* mutant was similar to the WT (Fig. 5A), although the plug domain was absent in the *sepN* mutant (Fig. 4). This is in line with the phenotype of the *sepN* mutant, which is different from the typical Δ*fraD* phenotype, like filament fragmentation, inability of diazotrophic growth and a reduced nanopore array. Proper localization of FraD in the *sepN* mutant challenges the assumption of FraD localizing to the plug module, since the latter was lacking in a *sepN* mutant. Instead of forming the plug module itself, FraD might rather be a linker connecting different SJ modules. Our data suggest an essential structural impact of SepN on the formation of the plug module. Assuming SepN as a SJ-protein, the lower fluorescence signal observed for SepN-sfGFP in the absence of FraD (Fig. 5B) could also arise from the reduced number of nanopores in the absence of *fraD*, which might lead to a fluorescence signal below the detection limit. It therefore remains unknown, if the localization of SepN is dependent on the presence of FraD.

In reverse co-IPs using GFP-SepN as bait, FraD was the most abundant protein. Also, FraC, which is encoded upstream of and in the same operon as FraD, was detected in the reverse co-IP with and without glutaraldehyde treatment among the twenty most abundant proteins. In contrast, FraC was never detected in co-IPs directed against FraD and also, direct protein-protein interaction was never shown for FraC and FraD. Hence, SepN may form a linker between FraC and FraD.

In conclusion, we identified SepN as an essential protein for the assembly of the SJ complex. It interacts with FraD and is most likely itself a structural SJ component. Moreover, the *sepN* mutant allowed to elucidate the so far elusive role of the SJ plug module as important for full inhibition of intercellular communication and hence gating behavior of the cell-cell communication structures of the *Nostoc* filament.

## Author contribution

AKK: designed and performed most of the experiments, analyzed and interpreted data, drafted the work and wrote the manuscript.

PT and GLW: designed and performed ECT experiments, analyzed and interpreted the corresponding data, revised the manuscript critically.

MP: designed and supervised the research in Zürich, interpreted data, revised the manuscript critically.

IM: designed and supervised the research in Tübingen, made considerable contributions to the design of the study, interpreted data, revised the manuscript critically.

All authors approved the final manuscript.

## Acknowledgment

We thank Mirita Franz-Wachtel of the Proteom Center Tübingen for performing LC-MS/MS, and Jan Bornikoel for construction of the C.K3t4 cassette. We thank Karl Forchhammer for fruitful discussions. Furthermore, we thank Teresa Müller for performing the FRAP of DR825.848 and Enrique Flores for providing strain CSVT2. Work in Tübingen was supported by the German research foundation (GRK1708 and DFG-MA1359/7). We acknowledge instrument access at the imaging platform ScopeM at ETH Zürich. Work in Zurich was supported by the Boehringer Ingelheim Fonds, the NOMIS foundation, and SNSF (31003A_179255).

