## supplemental table for "SepN is essential for assembly and gating of septal junctions in *Nostoc* sp. PCC 7120"

**Supplemental data:**

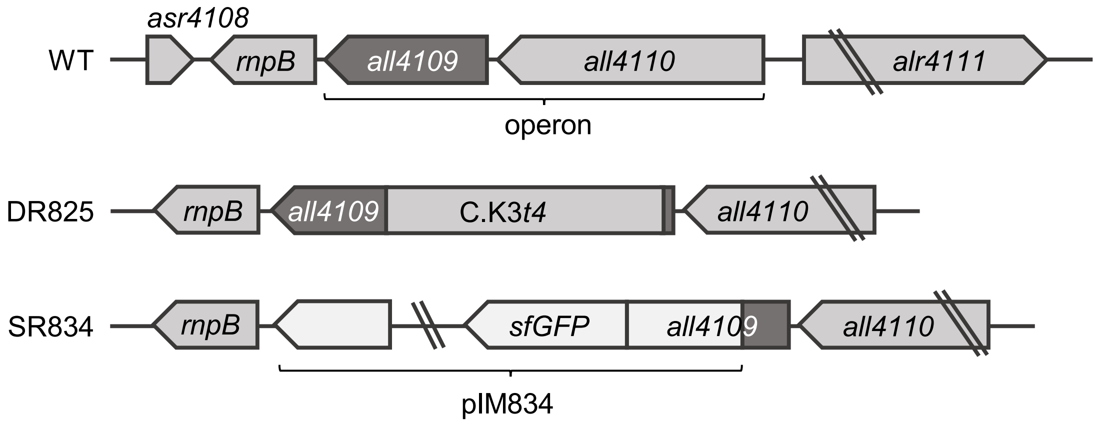


**Supplement Figure 1:** Schematic representation of the *all4109* genomic region.

For inactivation of *all4109* the neomycin resistance cassette C.K3*t*4 was inserted into the ORF of *all4109* via double homologous recombination creating strain DR825. Genomic *all4109* was exchanged for an *all4109-sfgfp* translation fusion via single homologous recombination with plasmid pIM834 yielding strain SR834.

**Supplement Table 1:** Strains and plasmids used in this work.

| ***Nostoc* sp. strains** | Relevant characteristics | Reference |
|---|---|---|
| PCC 7120 | wild type | Rippka *et al.,* 1979 |
| CSVT2 | Δ*fraD* (Δ*alr2393*) | Merino-Puerto *et al.,* 2010 |
| 7120.800 | $P_{fraCDE}$-*gfpmut2*, $Sm^r$, $Sp^r$ | This study |
| DR825 | *all4109*::C.K3*t4* cassette, $Nm^r$ | This study |
| SR834 | *all4109*-5xGS-*sfgfp*, $Sm^r$, $Sp^r$ | This study |
| DR825.848 | *all4109*::C.K3*t4*, $P_{all4110/4109}$-*all4109*, $Nm^r$, $Sm^r$, $Sp^r$ | This study |
| CSVT2.SR834 | Δ*fraD* (*alr2393*), *all4109*-5xGS-*sfgfp*, $Sm^r$, $Sp^r$ | This study |
| ***Escherichia coli* strains** | | |
| NEB 10β | *Δ(ara-leu) 7697 araD139 fhuA* Δ*lacX74 galK16 galE15 e14- Φ80*d*lacZ*Δ*M15 recA1 relA1 endA1 nupG rpsL* ($Str^R$) *rph spoT1* Δ*(mrr-hsdRMS-mcrBC)* | NEB *biolabs* |
| HB101 | F–, *thi*-1, *hsd*S20 ($r_B^-$, $m_B^-$), *sup*E44, *rec*A13, *ara*-14, *leu*B6, *pro*A2, *lac*Y1, *gal*K2, *rps*L20 ($str^r$), *xyl*-5, *mtl*-1 | Sambrook *et al*., 1989 |
| J53 (RP-4) | R+, *met*, *pro* (RP-4: *Ap*, *Tc*, *Km*, $Tra^+$, *IncP*) | Wolk *et al*., 1984 |
| **Plasmids** | | |
| pIM800 | $P_{fraCDE}$-*gfpmut2* in pRL1049, $Sm^r$, $Sp^r$ | This study |
| pIM825 | C.K3*t4* cassette flanked by upstream and C-terminal fragment of *all4109* in pRL277, $Sm^r$, $Sp^r$, $Km^r$ | This study |
| pIM834 | C-terminal *all4109*-fragment fused to 5xGS-linker and *sfgpf* in pRL277, $Sm^r$, $Sp^r$ | This study |
| pIM848 | $P_{all4110/4109}$-*all4109* in pRL1049, $Sm^r$, $Sp^r$ | This study |
| pIM779 | $P_{fraCDE}$-*gfpmut2-fraD* in pRL1049, $Sm^r$, $Sp^r$ | Weiss *et al*., 2019 |
| pIM660.2 | C-terminal *alr3353*-fragment fused to 5xGS-linker and *sfgfp* in pRL277, $Sm^r$, $Sp^r$ | Bornikoel, unpublished |
| pRL1049 | Self-replicating plasmid for *Anabaena* sp., $Sm^r$, $Sp^r$ | Black and Wolk, 1994 |
| pRL277 | Non-replicating, mobilizable vector containing *sacB*, $Sm^r$, $Sp^r$ | Black *et al*., 1993 |
| pRL528 | Helper plasmid for mobilization used in triparental mating, $Cm^r$ | Wolk *et al*., 1984 |

Sm: Streptomycin, Sp: Spectinomycin, Km: Kanamycin, Nm: Neomycin, Cm: Chloramphenicol

**Supplement Table 2:** Oligonucleotides used in this work.

| Oligo # | Sequence (5' → 3') |
|---|---|
| 1383 | TCTAGAGGATCTCAATGAATA |
| 1384 | ATGCTTGTAAACCGTTTTG |
| 1444 | TAGTGGATCCGGTAGTGGATCCGGTAGCGCATCAAAAGGTGAAGAATTATTTAC |
| 1445 | GCCAGTTAATAGTTTGCGCAACGTTGTTGCCATTGCTGCATTATTTATATAATTCAT CCATACCATG |
| 1998 | GATATCCCGCAAGAGGCCCTTTCGTCTTCAAGAATTCTGCCGTTCCTTGTCATCTG |
| 2235 | CCACAACGGTTTCCCTCTACCGGGATCCGGTTATTTGTATAGTTCATCCATGCCAT GTG |
| 2397 | ATTCATTGAGATCCTCTAGATGGTCAGTACTCCTAGTC |
| 2398 | CAAAACGGTTTACAAGCATATTGGACTTATGCCCTACC |
| 2399 | ATGGCAGAAATTCGATATCTAGATCTCGAGTGCTCGATGCGATTATTG |
| 2400 | TAATAGTTTGCGCAACGTTGTTGCCATTGCTGCAGGTTGTCAGTTGCCAGTTG |
| 2446 | GCTTTGCAGGCGTGAG |
| 2447 | TCGCACTGGACGTTATC |
| 2450 | ATGGCAGAAATTCGATATCTAGATCTCGATTGGTGTATCTTATATATT |
| 2451 | ACCGGATCCACTACCGGATCCACTACCCTTTTTATTCCAGGGTAGG |
| 2511 | AGAGGCCCTTTCGTCTTCAAGAATTTTGCCGTCAGGCTTAG |
| 2512 | TGGTCAGTACTCCTAGTCATGCAGTTATACCCGAAACTT |
| 2513 | ATGACTAGGAGTACTGACC |
| 2514 | GACCACAACGGTTTCCCTCTACCGGTTATTTCTTTTTATTCCAGGGTAG |

**Supplement Table 3**: Performed co-IPs with respective controls.

| Antibody | Co-IP | Sample | Control |
|---|---|---|---|
| α-GFP | 1 | CSVT2.779 | CSVT2.779 empty beads |
| | | CSVT2.779 | |
| | 2 | CSVT2.779 | 7120.800 |
| | | CSVT2.779 | |
| | 3 | CSVT2.779 | CSVT2.779 empty beads |
| α-FraD | 1 | WT | WT pre-immunserum |
| | 2 | | CSVT2 |
| | 3 | | |

- glutaraldehyde (white)
+ glutarladehyde (grey)

CSVT2.779: Δ*fraD* + p(*gfpmut2-fraD*); 7120.800: WT + p(*gfpmut2*); CSVT2: Δ*fraD*.
